# Redefining the Role of the EryM Acetyltransferase in Natural Product Biosynthetic Pathways

**DOI:** 10.1101/2025.03.07.642089

**Authors:** Yihua Li, Xunkun Liu, Natalia R. Harris, Jacquelyn R. Roberts, Estefanía Martínez Valdivia, Xinrui Ji, Janet L. Smith

**Affiliations:** Life Sciences Institute, University of Michigan, Ann Arbor, MI 48109 USA; Department of Biophysics, University of Michigan, Ann Arbor, MI 48109 USA; Department of Chemistry, University of Michigan, Ann Arbor, MI 48109 USA; Department of Biological Chemistry, University of Michigan, Ann Arbor, MI 48109 USA; Program in Chemical Biology, University of Michigan, Ann Arbor, MI 48109 USA; Genentech, South San Francisco, CA 94080 USA

## Abstract

The GNAT (GCN5-related *N*-acetyltransferase) superfamily comprises enzymes with a conserved fold and diverse catalytic activities, including primarily acyl transfer, with a few examples of decarboxylation. EryM, a GNAT enzyme from *Saccharopolyspora erythraea*, has been implicated in both erythromycin and erythrochelin biosynthesis, with dual functionality as an acetyltransferase and a decarboxylase. Despite an historical association with malonyl-coenzyme A decarboxylation activity, the structural basis for this dual activity has remained unknown as its close homologs were identified with only acyl transfer activity. Here, crystal structures of EryM in its free form (2.5 Å) and in complex with acetyl-CoA (2.9 Å) reveal insights into the active site architecture and substrate interactions. Functional assays demonstrate that EryM catalyzes acyl transfer but lacks decarboxylation activity, challenging long-standing assumptions about its biosynthetic role. Comparative analysis of EryM and homologs in siderophore biosynthetic pathways highlights a conserved catalytic pocket with an essential His and identically positioned side chains common to GNAT enzymes for *N*-acyl transfer from CoA to primary hydroxylamine substrates. Bioinformatic analysis defines a large GNAT subfamily that is broadly distributed in the microbial world. These findings redefine EryM as an acetyltransferase and provide a foundation for understanding GNAT functional diversity in natural product biosynthesis.

## Introduction

Many enzyme superfamilies include members adapted to diverse catalytic and non-catalytic functions. The GNAT (GCN5-related *N*-acetyltransferase) superfamily is an outstanding example^1,2^. Most characterized GNAT proteins transfer an acyl group from coenzyme A (CoA) to a nitrogen atom on a protein or small-molecule substrate^1,3^. The founding member and the most extensively studied GNATs are histone lysine acetyltransferases (HATs), which exist in four GNAT subfamilies^4^. Other *N*-acyltransferase GNATs act on a variety of protein and small-molecule substrates, notably the aminoglycosides^1^. Within the common GNAT fold, the active site is located in a V-shaped cleft near the center of a curved β-sheet^2^. Sequence identity is low across the superfamily, and the active sites of *N*-acyltransferase GNATs lack conserved amino acids, although the general base for deprotonation of the nitrogen acceptor has been identified or proposed for some^1–5^.

A few GNAT proteins do not catalyze acyl transfer, but instead decarboxylate acyl groups on CoA or acyl carrier protein (ACP) substrates. The unexpected decarboxylase activity was initially characterized for the GNAT domain within CurA from a type I modular polyketide synthase (PKS)^6^. The PKS GNAT decarboxylases exist in multi-enzyme proteins of pathways for curacin A, saxitoxin and gephyronic acid where they generate starter units by decarboxylation of malonyl-, methylmalonyl- or dimethylmalonyl-ACP, respectively^7^. Subsequent to the discovery of the PKS GNAT decarboxylases, the GNAT fold was discovered in several bacterial and human malonyl-CoA (Mal-CoA) decarboxylases^8^. All of these GNAT decarboxylases possess conserved His and Ser/Thr amino acids, which were shown to be essential for the decarboxylase activity of the CurA GNAT^6^. As in the active sites of the GNAT *N*-acyltransferases, the His and Ser/Thr amino acids are located in the V-shaped cleft^6^ although the enzymes lack acyl transfer activity^7^. We designate these decarboxylases as the CurA subfamily within the GNAT superfamily.

In a recent study to characterize GNAT decarboxylases in PKS starter modules^7^, we identified EryM, a GNAT with a storied history and apparently dual acyltransferase and decarboxylase functions^9–13^. In 1984, Kolattukudy and co-workers isolated a putative Mal-CoA decarboxylase from the erythromycin producer *S. erythraea* and proposed a role in erythromycin biosynthesis based on the strong correlation of antibiotic production with levels of protein and activity^9^. In 1994, they identified the gene and showed that its disruption substantially reduced erythromycin production^10^. Addition of propionate to the growth medium of the knock-out strain restored erythromycin production, so the gene was named *eryM*. The biosynthetic logic for the erythromycin macrolactone had been established previously^14^, but the source of the propionyl-CoA (Prop-CoA) starter unit was unknown. Thus, EryM was proposed to provide propionate by decarboxylation of methylmalonyl-CoA (MeMal-CoA)^10^ (Figure 1A), although this has not been shown directly. In 2007, EryM acquired another name, SACE_1304, when Leadlay and co-workers reported the *S. erythraea* genome sequence^15^. Of the 417 amino acids in EryM/SACE_1304, the C-terminal ∼185 residues constitute a GNAT domain.

**Figure 1.**
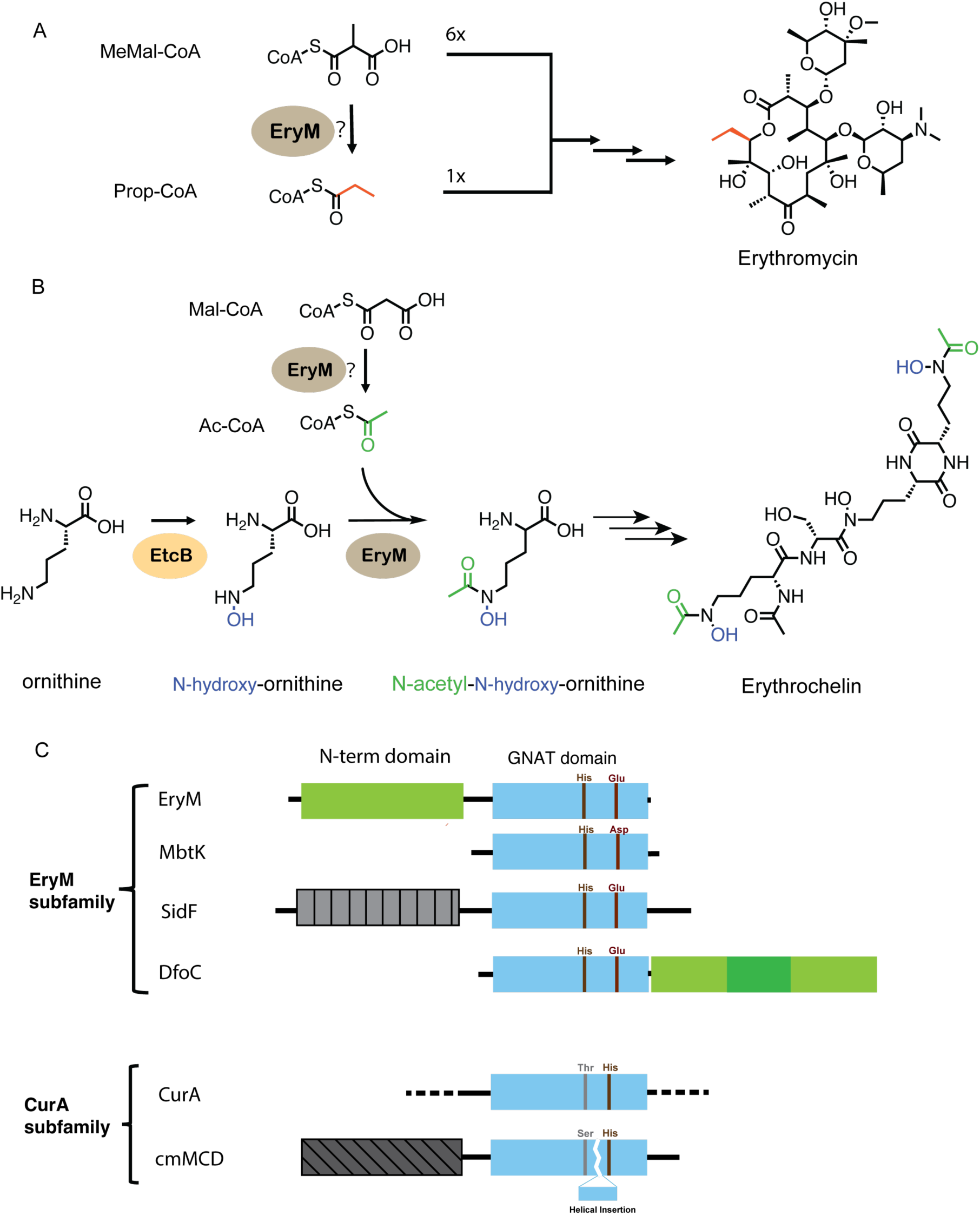
EryM functions and domain structure. **A**. Erythromycin biosynthesis. EryM is proposed to form the starter unit, Prop-CoA, by decarboxylation of MeMal-CoA^10^. **B**. Erythrochelin biosynthesis. In addition to proposed decarboxylation of Mal-CoA, EryM acetylated hOrn after Orn hydroxylation by EtcB^11,13^. **C**. EryM and CurA GNAT subfamilies showing the different locations of highly conserved amino acids. These residues are essential to the decarboxylation activity of CurA^6^ and the acyl transfer activity of MbtK^24^.

In 2010-2011, a second function was identified for EryM when the Marahiel and Leadlay groups reported the biosynthesis of the siderophore erythrochelin by *S. erythraea*^11–13^ (Figure 1B). The initial steps are hydroxylation of ornithine (Orn) by EtcB (flavin-dependent δ-*N*-ornithine monooxygenase also known as ErcB and SACE_3033) to form δ-*N*-hydroxy-ornithine (hOrn), then acetylation by EryM to form δ*-N*-acetyl-δ*-N*-hydroxy-ornithine (Ac-hOrn)^11,12^. Both groups also proposed that EryM produces acetyl-CoA (Ac-CoA) by decarboxylation of Mal-CoA^11,13^. Ac-hOrn then serves as a building block for the nonribosomal peptide synthetase (NRPS) EtcD/ErcD/SACE_3035 (Figure 1B). Consistent with roles in the biosynthesis of both erythromycin and erythrochelin, the location of *eryM* in the *S. erythraea* genome is remote from the biosynthetic gene clusters (BGCs) of both natural products^11,15^.

Several proteins with GNAT domains have similar sequences to the EryM GNAT domain and are annotated as acyltransferases in the biosynthesis of siderophores including aerobactin (IucB in a bacterial NRPS-independent siderophore (NIS) pathway)^16^, mycobactin (MbtK in a bacterial hybrid NRPS-PKS pathway)^17^, desferrioxamine E (DfoC in a bacterial NIS pathway)^18^, as well as fusarinine and ferrichrome (SidF and SidL in a fungal NRPS pathway)^19–21^. In crystal structures of MbtK^22^, DfoC^23^ and SidF^21^, conserved His and Glu amino acids are located near the prototypical GNAT active site, and are also conserved in EryM, IucB and SidL. Thus we designate these proteins as the EryM GNAT subfamily (Figure 1C). However, despite their sequence similarity to the EryM GNAT domain, no decarboxylase activity has been reported or proposed for MbtK, SidF, SidL, DfoC or IucB, leaving open the question of what features of the EryM structure enable its dual activity. These proteins have variable domain organization, including GNAT-only (MbtK, IucB), N-terminal GNAT (DfoC), and C-terminal GNAT (EryM, SidF, SidL) (Figure 1C). The EryM N-terminal domain has no known function or obvious ancestry, differs from the related SidF and SidL N-terminal domains, and thus may contribute to any EryM dual activity (Figure 1C).

At the sequence level, the EryM and CurA GNAT subfamilies are distinct from one another with highly conserved amino acids at different positions in their sequences (Figure 1C). The domains beyond the GNAT are dissimilar. In the CurA GNAT subfamily, the GNAT domains of prokaryotic and eukaryotic Mal-CoA decarboxylases are preceded by N-terminal domains that are unrelated to the EryM N-terminal domain, and the PKS GNAT decarboxylases lack N-terminal domains. Interestingly, the invariant amino acids essential for activity of the CurA GNAT decarboxylase subfamily cluster on the opposite side of the central GNAT β-sheet from the conserved His and Glu residues in the EryM subfamily^6^. Thus, the GNAT superfamily may have evolved yet another active site architecture to adapt to dual activity in EryM. Intrigued by this observation, we investigated the EryM structure and its proposed catalytic functions in erythromycin and erythrochelin biosynthesis. Here we report crystal structures of EryM and the results of functional assays revealing that EryM catalyzes acyl transfer but not decarboxylation.

## Results

### The structure of EryM

To inform our investigation of EryM catalytic function, we solved crystal structures of the EryM free protein (2.5 Å) and a binary complex with Ac-CoA (2.9 Å, Figure 2, Table S1). EryM is a dimer in the crystal structures, as in solution (Figure S1A, B). The dimer interface is formed by the N-terminal domain (amino acids 1-230). The peripheral C-terminal domain (amino acids 233-417) has the classic GNAT fold.

**Figure 2.**
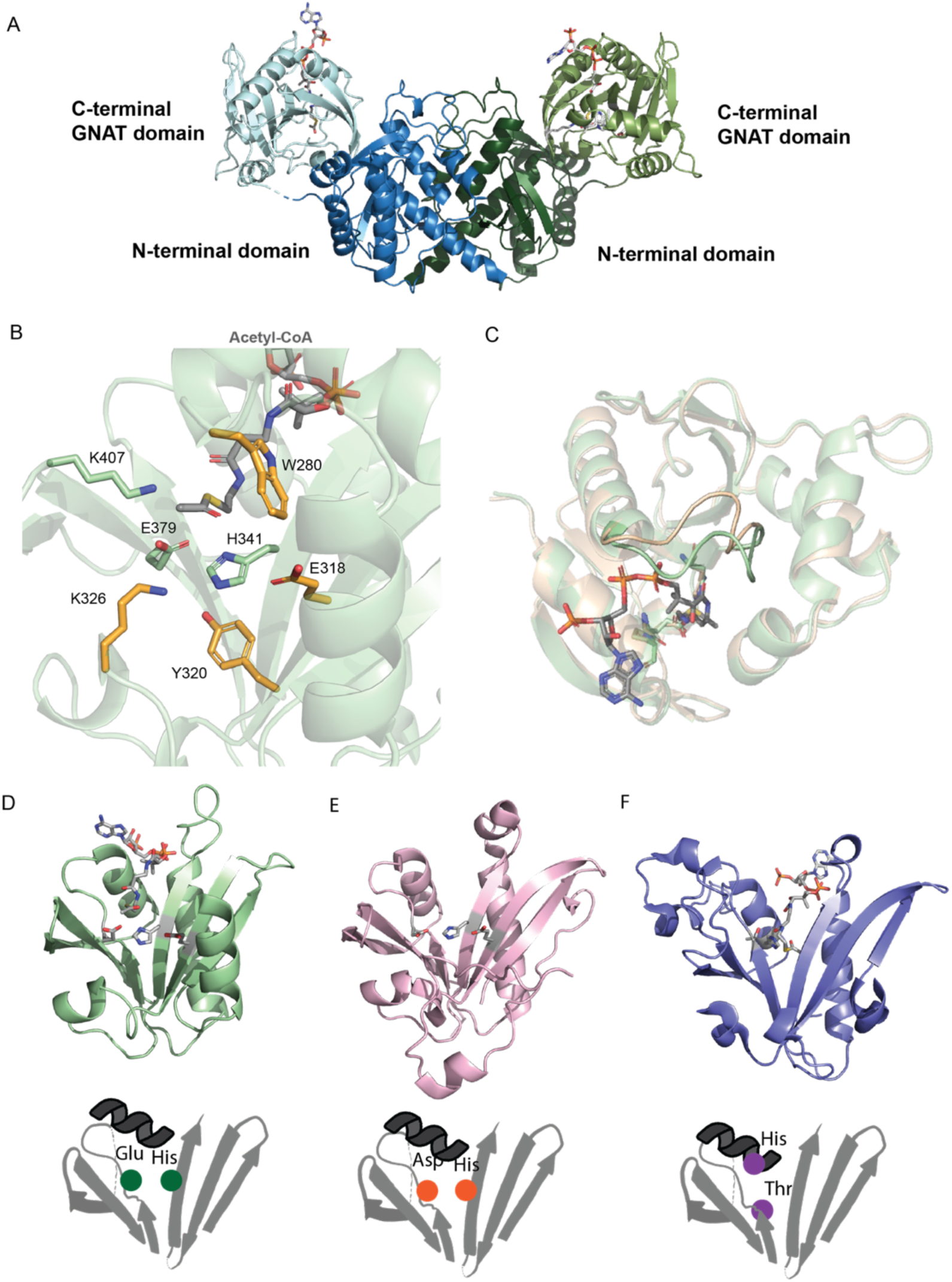
Structure of EryM and comparison with homologs. **A**. EryM dimer in complex with Ac-CoA. Subunits are in blue and green, each with a darker shade for the N-terminal domain and lighter for the GNAT domain. **B**. Ac-CoA binding pocket with nearby side chains in stick form. His341, Glu379 and Lys407 (green C atoms) are within 4 Å of the thioester of Ac-CoA (gray C), and Tyr320, Glu318, Lys326 and Trp280 (orange C) are within 4 Å of His341. **C**. Superposition of EryM Ac-CoA complex (tan) and the free protein (green). **D**. EryM GNAT domain with bound Ac-CoA and Glu318, His341 and Glu379 side chains in stick form. The illustration below shows these residues on the front side of the β-sheet. **E**. MbtK^22^ (PDB 1YK3) with Glu107, His130 and Asp168 side chains in stick form, positioned identically to the analogous residues in EryM. **F**. CurA GNAT domain^6^ (PDB 2REF) showing His389 and Thr355 side chains (stick form) on the back side of the β-sheet.

The EryM N-terminal domain has undetectable sequence similarity to any domain in the conserved domain database^25^. However, the topology is similar to a portion of an adenylating ligase in several siderophore biosynthetic pathways, including those for achromobactin (AcsD)^26^, petrobactin (AsbB)^27^, aerobactin (IucA)^28^ and desferrioxamine E (DfoC C-terminal domain)^23^. These ligases possess thumb, palm and fingers subdomains^26^, but the EryM N-terminal domain includes only the thumb and thumb-adjacent regions of the palm, representing internal deletions of ∼360 amino acids in the ∼600-residue ligase. EryM thus lacks an ATP binding site, but has an electropositive cleft between the vestigial thumb and palm subdomains (Figure S2A, B, C). Notably, in the *iuc* BGC for aerobactin biosynthesis, *iucB* encodes an acetyltransferase homolog of the EryM C-terminal GNAT domain (Figure S2D)^16^. Thus, the *iucA*-*iucB* and *eryM* genes may share a common ancestor, and *eryM* may be the product of deletion and fusion events in which the deletion led to an EryM with a non-functional ligase vestige. The EryM dimer interface (Figure S2) aligns with helices of IucA, but this region is not part of any subunit interface in the tetrameric IucA.

The EryM C-terminal GNAT domain has the characteristic central seven-stranded β-sheet flanked by α-helices with V-shaped cleft created by splaying apart of β-strands in the center of the β-sheet (Figure 2). Ac-CoA binds to EryM with the pantetheine arm resting in the V-shaped cleft (Figure 2B), as in structures of other GNATs^1^. The acetyl thioester is surrounded by conserved residues His341, Glu379, Lys407, and Trp280 (Figure 2B, S3). The CoA diphosphate binds to the backbone amides of Gly residues at the start of an α-helix (Figure 2C). A shift of the loop preceding this helix is the only structural difference between the Ac-CoA complex and free EryM. The side chains of His341 and Glu379 in the V-shaped cleft point towards the Ac-CoA thioester (Figure 2D). Glu318 and His341 are invariant and Glu379 is a conserved carboxylate in the EryM GNAT subfamily. Acyl transfer activity has been demonstrated for EryM and three of these homologs, all from siderophore biosynthetic pathways (Figure S4). MbtK/Rv1347 from *Mycobacterium tuberculosis* transfers a fatty acyl chain from CoA to a hydroxylated lysine^17^; His130 and Asp168 in the V-shaped cleft (analogous to EryM His341 and Glu379) are essential for this activity^24^. The fungal SidF and SidL exhibit similar *N*-acyl transfer activities and possess conserved His and Glu residues ^20,21^ (Figure 1C, S4). The core GNAT structures of MbtK^22^, SidF^21^, DfoC^23^ and an uncharacterized bacterial gene product (PDB 2QML) are remarkably similar to the EryM GNAT (Cα RMSD = 1.0 – 1.3 Å) despite their modest sequence identity (23%-36% identity to EryM). A number of conserved amino acids surrounding the acyl-CoA pocket superimpose exactly in these structures, including EryM Trp280, Glu318, Tyr320, Asp325, His341, Glu379 and Lys407 (Figure S3C,D).

In contrast to the conserved amino acids in EryM and these acyl transfer enzymes, the CurA GNAT decarboxylase subfamily has conserved His and Ser/Thr located in the V-shaped cleft but on the opposite side of the β-sheet from the conserved His and Glu/Asp in EryM (Fig 2D,E). In CurA (Figure 2F), His389 and Thr355 are essential for decarboxylation activity^6^. The analogous positions in EryM are nonconserved Phe391 and invariant Pro380. Therefore, if EryM possesses decarboxylation activity, it must employ different amino acids than these decarboxylases, potentially the invariant functional groups of Glu318, His341 or Glu379.

### Evolutionary Insights and Homologs of EryM

To understand the biological distribution of EryM and search for any homologs of known function, we sought to identify all protein sequences containing both N-terminal and GNAT domains homologous to those of EryM. Non-redundant protein sequences in the RefSeq translatome (genomic, metagenomic and transcriptomic data)^29^ were probed with the full-length EryM sequence. We then used an in-house script^30^ to eliminate hits that did not align with both EryM domains and to extract the EryM-aligned regions from hits within larger multi-domain proteins. This yielded a total of only 58 unique sequences, all from the *Pseudonocardiaceae* family (Actinomycetes bacteria) and none that reside within a larger multi-domain protein. None of the hits is a previously characterized protein, and none has the His and Ser/Thr amino acids essential for decarboxylase activity in the CurA GNAT decarboxylase subfamily. The 58 sequences are highly similar, indicative of a common function. Sequence identity is especially high in the GNAT domain, where the average pairwise identity is 72% (minimum 37%, median 74%). All identified sequences share the conserved His (His341) and carboxylate (Glu318 and Glu379) residues that cluster around the Ac-CoA thioester in the EryM structure (Figure 2B). These amino acids are located within conserved sequence islands and represent motifs of the EryM GNAT subfamily: the DxGxH-G on β-strand D containing His341, the YxExY on β-strand C containing Glu318, and VVAE(D)P on β-strand E containing EryM Glu379 (Figure S3D). While they are less conserved than the GNAT domains, the 58 N-terminal domains are also highly similar (58% average pairwise identity, minimum 25%, median 55%). Among the few highly conserved amino acids in the N-terminal domain, the His27 and Arg91 side chains form hydrogen bonds with the GNAT domain. The limited biological distribution and novel domain organization consisting of a vestigial ligase fragment fused to a GNAT domain suggests that recent deletion and gene fusion events occurred within the *Pseudonocardiaceae* family.

In contrast to the few sequences that are homologs of full-length EryM, including both N-terminal and GNAT domains, we identified thousands of homologs with conservation of Glu318, His341 and Glu379 when probing with the GNAT domain alone. For this analysis, we searched the ClusteredNR database^31^, where each cluster consists of sequences with greater than 90% identity to one another and within 90% of the length of longest member. A total of 8,757 sequence clusters were identified. They are distributed among bacteria (77%), fungi (22%) and archaea (1%), but not elsewhere, underscoring the widespread distribution of the EryM GNAT subfamily across diverse microbial species.

### The Function of EryM

The ability of EryM to decarboxylate CoA species was not assayed directly in the published studies of erythrochelin biosynthesis^11,13^. Moreover, the active site location (Fig. 2D,E,F), structure (Figure 2B) and conserved amino acids in the EryM GNAT subfamily differ from those of the CurA GNAT decarboxylase subfamily. Therefore, we first tested whether EryM could decarboxylate Mal-CoA or MeMal-CoA by incubating each CoA species or its non-carboxylated analog, Ac-CoA or Prop-CoA, with or without EryM, under conditions that included those previously published^13^. CoA species were separated by high-pressure liquid chromatography (HPLC), detected by absorbance at 258 nm (A_258_) and mass spectrometry (MS), and quantitated by A_258_. In these experiments, it was important to control for any spontaneous decarboxylation or de-acylation during the incubation period as well as any contamination from decarboxylated or de-acylated species in (even freshly prepared) CoA stocks (Figure S5). The concentrations of the carboxylated CoAs, Mal-CoA or MeMal-CoA, were unaltered by EryM, although we detected a low level of de-acylation of Ac-CoA and Prop-CoA in the presence of EryM (Figure 3). Thus, in these conditions, EryM was not a decarboxylase.

**Figure 3.**
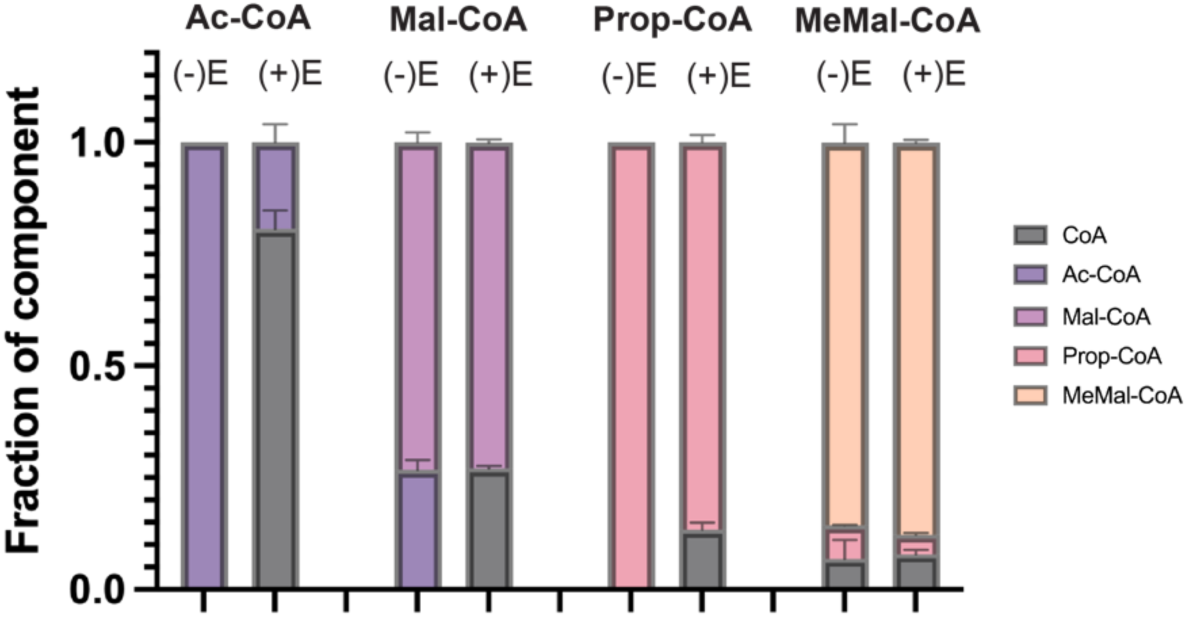
Decarboxylation analysis of CoA derivatives in the presence or absence of EryM. Ac-CoA, Mal-CoA, Prop-CoA or MeMal-CoA were incubated with or without EryM, and substrates/products were separated and detected by absorbance at 258 nm (A_258_). The Mal-CoA stock had ∼25% Ac-CoA contamination, which was de-acetylated during the 1-hr incubation (Figure S5). EryM had no detectable decarboxylation activity with either Mal-CoA or MeMal-CoA.

In erythrochelin biosynthesis, EryM was shown to acetylate hOrn^11,13^. Thus, we tested the four CoA species for acyl transfer to hOrn, using LC-MS to detect ornithine species, again using published reaction conditions^13^. Under the assay conditions (1 hr, 25°C), EryM fully converted hOrn to δ-acetyl-δ-*N*-hydroxyornithine (Ac-hOrn) with Ac-CoA as an acetyl donor (Figure 4A). With a Mal-CoA donor, Ac-hOrn was formed at much lower levels that were attributable to Ac-CoA contamination of the Mal-CoA stock (Figure S5). Interestingly, Prop-CoA was also an effective acyl donor; EryM fully converted hOrn to δ-*N*-propionyl-hOrn (Prop-hOrn). Like Mal-CoA, MeMal-CoA yielded low levels of Prop-hOrn, again attributable to Prop-CoA contaminantion of the MeMal-CoA stock (Figure S5). No malonyl-hOrn or methylmalonyl-hOrn was detected in these experiments. Thus we conclude that EryM forms acyl-hOrn from non-carboxylated CoAs, but does not decarboxylate Mal-CoA or MeMal-CoA before, during or after acyl transfer.

**Figure 4.**
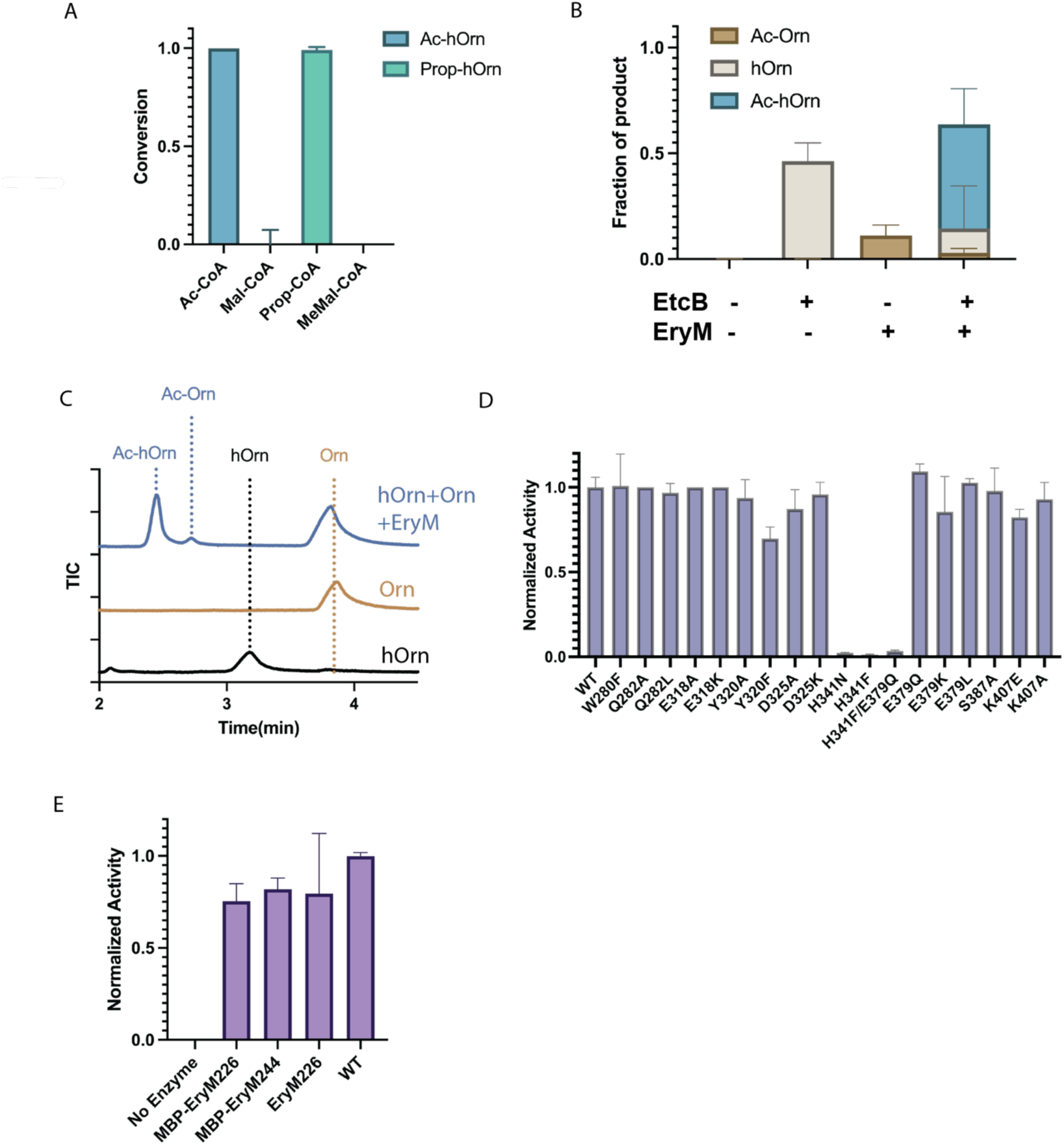
EryM acyl transfer activity. **A**. Conversion of acyl-CoAs to acyl-hOrn species by EryM in a 1-hr reaction. Data are corrected for Ac-CoA contamination of Mal-CoA stocks and Prop-CoA contamination of MeMal-CoA. **B**. Coupled assay of EryM and EtcB, using an Ac-CoA donor. **C.** Formation of acetylation products in the presence of both hOrn and Orn. **D.** Normalized acetyl transfer activity of EryM variants. **E.** Normalized acetyl transfer activity of truncated EryM GNAT domain variants. Error bars represent the variance of triplicate experiments.

The hOrn acceptor substrate is the product of ornithine hydroxylation by EtcB^11,13^. To examine the acetyltransferase activity of EryM for hOrn, we purified recombinant EtcB, solved a crystal structure (Table S1, Figure S1C, D), and used the enzyme in a coupled assay with EryM by incubating Orn, Ac-CoA with or without EryM and/or EtcB and its co-factors FAD and NADPH. After 4 hr at 30°C, the reaction mixtures were quenched and analyzed by LC-MS. We detected the formation of hOrn by EtcB irrespective of EryM presence, and the formation of Ac-hOrn in the presence of both EryM and EtcB (Figure 4B). EryM acted on both acetyl group acceptors, generating Ac-Orn from Orn and Ac-hOrn from hOrn. To determine the substrate preference, we conducted a direct competition assay by incubating EryM with Ac-CoA and an equimolar mixture of hOrn and Orn (Figure 4C). Under these conditions, EryM showed a strong preference for hOrn, producing Ac-hOrn at levels approximately 11-fold greater than Ac-Orn.

Amino acids within ∼4 Å of the Ac-CoA thioester (Figure 2B) were probed by site-directed mutagenesis and assays with Ac-CoA and hOrn substrates (Figure 4D). Compared to wild type EryM, substitutions at His341 (H341F, H341N) led to nearly undetectable acetyl transfer activity. In contrast, substitutions for conserved Trp280 (W280F), Glu318 (E318K, E318A), Glu379 (E379Q, E379K, E379L) or Lys407 (K407E, K407A) had little impact on activity. These results suggest that His341 may serve as the general base for deprotonation of the acceptor substrate hOrn. Substitutions for other conserved amino acids near the substrate thioester or His341, Gln282 (Q282L, Q282A), Tyr320 (Y320F, Y320A), Asp325 (D325K, D325A), or Ser387 (S387A), also had no effect on the EryM ability to acetylate hOrn.

To investigate whether the N-terminal domain has a role in EryM catalysis, we excised the C-terminal GNAT domain and purified fragments with (EryM226, residues 226-417) or without (EryM244, 244-417) the inter-domain linker. EryM226 and EryM244 had marginal stability, suggesting the natural EryM N-terminal domain serves as a stabilizing scaffold. Both of the EryM fragments were stabilized by N-terminal fusion with the maltose binding protein (MBP). Proteolytic cleavage of the MBP tag led to aggregation, and only EryM226 yielded sufficient quantities for assay. In assays with Ac-CoA and hOrn substrates, MBP-EryM244, MBP-EryM226 and EryM226 had near wild-type acetyltransferase activity, indicating that the N-terminal domain has no role in acetyl transfer (Figure 4E). A similar observation has been reported for SidF, where the N-terminal domain is a truncated GNAT domain that serves only as a scaffold^21^.

## Discussion

This study reveals that the EryM catalytic activity resides exclusively in the C-terminal GNAT domain (Figure 4E). The N-terminal domain is a fragment of an ancestral ligase and forms the subunit interface of the dimeric protein (Figures 2, S2). The two-domain architecture of EryM is unique to the *Pseudocardiaceae* family with fewer than 60 examples in the sequence database. We show that in addition to the previously demonstrated *N*-acetyltransferase activity^11,13^, EryM can also act on a Prop-CoA donor (Figure 4A). Both hOrn and Orn were acetyl acceptors, but EryM had a tenfold preference for the natural acceptor hOrn (Figure 4C). In a coupled assay, the hydroxylating oxygenase EtcB/ErcB produced hOrn from Orn for EryM acetylation (Figure 4B). EryM lacked any detectable decarboxylase activity (Figure 3).

### EryM GNAT structure and activity are like acyltransferases in other siderophore pathways

The EryM acyltransferase activity is like that of other members of the EryM GNAT subfamily that act in siderophore pathways. MbtK, IucB, SidF, SidL and DfoC all transfer acyl groups from CoA to a hydroxylated primary amine – δ-*N*-hydroxy-ornithine (EryM, SidF^21^, SidL^20^), ε-*N*-hydroxy-lysine (MbtK^17^, IucB^16^), or *N*-hydroxy-cadaverine (DfoC^23^). The natural acyl-CoA substrates vary considerably – acetyl (EryM, IucB, SidF, SidL), C_2-19_ acyl (MbtK^32^), anhydromevalonyl (SidF^33^), or succinyl (DfoC). In all cases tested, the nitrogen acceptor substrate is first hydroxylated, then acylated, and finally incorporated into the siderophore skeleton.

Within the EryM GNAT subfamily, the active site structure is highly conserved with His and Glu/Asp side chains near the acyl-CoA thioester. Five other conserved amino acids (EryM Trp280, Glu318, Tyr320, Asp325, Lys407) occupy identical positions in all structures (Figure S3C). EryM His341 is the only of these amino acids that is essential for acetyl transfer activity (Figure 4D), consistent with results for the MbtK homolog^24^. This is a remarkable level of conservation given the modest sequence identity among these enzymes (23-36% identity to EryM). This active-site conservation likely derives from the common activity of acyl transfer to the nitrogen of a primary *N*-hydroxylamine. The enzymes must position the acceptor substrate precisely in order for deprotonation and acyl transfer to occur at the nitrogen and not the oxygen. We attempted to understand the structural basis for nitrogen-selective acylation with numerous efforts to crystallize an hOrn complex of EryM, but these were unsuccessful. While the active site and CoA-binding pocket are highly conserved, structural differences in the outer regions of the GNAT likely contribute to substrate selectivity and affinity. In SidF^21^ and MbtK^22^, for example, a loop near the entrance to the presumed amino acid-binding pocket may obstruct or regulate access for larger substrates. Additionally, a hydrophobic pocket near the thioester site of MbtK can accommodate long acyl chains, but is absent from EryM and SidF. This structural feature aligns with the MbtK role to transfer acyl chains as long as C_20_ in mycobactin synthesis.

### EryM is not a decarboxylase

Most of our assay results mirror those of previous studies^11–13^, but we detected no decarboxylase activity for EryM in direct assays where CoA levels were followed (Figure 3). Assays were performed under a wide range of conditions, including those published^13^. Based on our experience with the instability of carboxylated CoAs^34^, we accounted for nonenzymatic decarboxylation of Mal-CoA and MeMal-CoA under the long assay times as well as decarboxylated and deacylated contaminants in freshly prepared solutions (Figure S5). We also considered that decarboxylation and acetylation may be coupled. However, even in the presence of the hOrn acetyl acceptor, we detected no decarboxylation of either Mal-CoA or MeMal-CoA (Figure 4A). Additionally, the crystal structure revealed that the EryM N-terminal domain is a vestigial ligase fragment (Figure S2) and is unlike any known decarboxylase. Thus we conclude that EryM is not a decarboxylase and not the proximal source of either Prop-CoA for erythromycin biosynthesis or Ac-CoA for erythrochelin biosynthesis.

Our study was intended to characterize the EryM dual functions of decarboxylation and acyl transfer and to discover its structural basis. We hypothesized that both activities resided in the GNAT domain, although others had hinted that the N-terminal domain may be a decarboxylase. The results disproved both hypotheses: EryM is not a decarboxylase and the N-terminal domain has no catalytic function. Consistent with these findings, the adaptation of a catalytic domain to non-catalytic function occurs in other natural product biosynthetic pathways. For example, truncated GNAT domains exist at the N-terminus of SidF^21^ and in the starting module of the PKS-NRPS pathway for apratoxin A^35^. Similarly, unused catalytic domains exist in the curacin A and jamaicamide pathways^36^. These results illustrate the facile microbial exchange of DNA and rapid evolution of natural product biosynthetic pathways.

### EryM does not provide Prop-CoA for erythromycin biosynthesis

All of the characterized EryM GNAT subfamily members, including EryM, IucB, SidF, SidL, MbtK or DfoC, function as acyltransferases, but none has been shown to decarboxylate Mal-CoA. Given that EryM is not a decarboxylase and *eryM* disruption blocks erythromycin production by *S. erythraea*^10^, its involvement in the pathway must be indirect. For example, two cytochrome P450 enzymes, EryF and EryK, catalyze late-stage hydroxylation reactions on the erythromycin macrolactone precursor, 6-deoxyerythronolide B (6-DEB)^37–39^ and may depend on erythrochelin to provide iron. Alternatively, EryM may be essential for another metabolic step leading to Prop-CoA. The location of *eryM* outside the *nrps5*/*erc* BGC for erythrochelin suggests that EryM does not act exclusively in the erythrochelin pathway, but catalyzes one or more other acyl transfer reactions in *S. erythraea*.

While the source of Prop-CoA for erythromycin biosynthesis is unknown, several lines of evidence suggest that branched-chain amino acid catabolism is the major source of Prop-CoA in *S. erythraea*^40^. Valine has been implicated as a precursor to the building blocks of macrolactones^41–43^, and is on-pathway to propionate^44^. Manipulation of either the *S. erythraea bkd* operon for degradation of branched-chain amino acids or its regulatory gene *pccD* significantly impacted erythromycin production^40^. Prop-CoA and MeMal-CoA are the building blocks for the erythromycin macrolactone, 6-DEB. A Prop-CoA carboxylase produces the MeMal-CoA extender unit directly from Prop-CoA, so it is difficult to disentangle the effects of genes on starter-unit vs. extender-unit availability. *S. erythraea* encodes several biotin-dependent acyl-CoA carboxylases^15,45^. Two of these, the mutli-domain SACE_4237^45^ and the multi-subunit SACE_3398-3400^46^, have been shown to produce MeMal-CoA from Prop-CoA and to affect intracellular Prop-CoA levels and erythromycin production. *S. erythraea* also encodes several propionyl-CoA synthetases, which convert propionate to Prop-CoA (SACE_0337, SACE_4729, SACE_3848, and SACE_1780). Among these, SACE_1780 overexpression resulted in increased erythromycin production^47^, consistent with the observation that propionate feeding overcame *eryM* disruption^10^. Together, these findings suggest that a pathway involving amino acid degradation generates Prop-CoA and MeMal-CoA for erythromycin biosynthesis.

In summary, through structural and biochemical analyses, we redefine EryM as an acetyltransferase and not a decarboxylase, clarifying its role in erythrochelin biosynthesis. The crystal structure of EryM with bound Ac-CoA shows how the enzyme interacts with its CoA donor substrate and reveals key features of the active site of the EryM GNAT subfamily. Comparative analysis with close homologs defines the EryM GNAT subfamily as catalysts for *N*-acyl transfer to primary hydroxylamine acceptor substrates and identifies conserved motifs for this activity.

## Acknowledgment

We thank Wenqing Feng for advice on LC/MS and David Akey for assistance with crystallographic analysis. This work was supported by NIH grant R01 DK042303 and the Rita Willis Professorship in the Life Sciences to JLS. GM/CA@APS is supported by NIH grant P30 GM138396 and the National Cancer Institute (ACB-12002).

## Author Contributions

YL and JLS designed the research. YL, XL, NRH, JRR, EMV and XJ performed the research. YL and JLS analyzed the data. YL and JLS wrote the paper with input from all authors.

## Method

### Construct Design

DNA encoding EryM (NCBI WP_011873320) and EtcB (NCBI WP_011873915) were amplified from synthetic genes (IDT and Twist Bioscience, respectively) with codon usage optimized for expression in *Escherichia coli*. The genes were inserted into the pMCSG7^48^ vector by Gibson assembly^49^ to create expression plasmids pEryM and pEtcB. Site-directed mutagenesis was done with the QuickChange Site-Directed Mutagenesis protocol (Agilent).

### Gene Expression and Protein Purification

*E. coli* BL21(DE3) cells were transformed with pEryM or pEtcB, encoding the protein of interest fused to an N-terminal His_6_ tag and a peptide cleavable by tobacco etch virus (TEV) protease. Cultures were grown at 37°C in 0.5 L TB with 100 μg/mL ampicillin to OD_600_ of 0.8, incubated1 hr at 20°C, induced with 0.2 mM IPTG, and grown 18 h. Cells were harvested by centrifugation at 8000 rpm for 30 min, resuspended in 40 mL buffer A (50 mM Tris pH 7.9, 300 mM NaCl, 10% (v/v) glycerol, 20 mM imidazole) with 0.1 mg/mL lysozyme, 0.05 mg/mL DNase, and 2 mM MgCl_2_, vortexed on ice for 30 min, and lysed by sonication on ice (30s with 30s intervals, repeat three times). Cell debris was removed by centrifugation at 38,760g for 30 min at 4°C. The supernatant was filtered and loaded onto a 5 mL HisTrap column (Cytiva). The column was washed with 50 mL buffer A and bound protein was eluted with a 50-mL gradient of 20-400 mM imidazole in buffer A. The eluted protein was concentrated to 5 mL and further purified via gel filtration chromatography (HiLoad Superdex S200, Cytiva) with buffer B (50 mM Tris pH 7.9, 150 mM NaCl, 10% (v/v) glycerol). The purified EryM was concentrated to ∼10 mg/mL and stored in aliquots at -80°C. Selenomethionyl (SeMet) EryM was produced by growing cultures in 0.5 L synthetic M9 minimal media supplemented with 100 μg/mL ampicillin and amino acids, except L-methionine was replaced with 0.04 mg/mL SeMet (Calibre Scientific UK). Culture growth and SeMet EryM purification protocols were identical to protocols for the wild-type. EtcB was prepared identically to wild-type EryM.

### Enzyme Assays and LC-MS Analysis

To test the decarboxylation activity, reaction mixtures (20 μL) containing 20 μM EryM, 1 mM acyl-CoA, 20 mM HEPES pH 8.0 and 100 mM NaCl were incubated at 25°C for 1 h. Conditions in which decarboxylation activity has been reported^13^ were also tested (4-8 hr incubation at 25°C in 20 mM TRIS pH 7.2, 100 mM NaCl). Reactions were quenched and protein precipitated by addition of 60 μL methanol. After centrifuging at 14,000 rpm for 10 min, reaction mixtures were subjected to analysis by liquid chromatography/mass spectrometry (LC/MS) using an Agilent Q-TOF 6545. Each quenched reaction mixture (5 μL) was loaded onto a high-performance liquid chromatography (HPLC) column (Agilent Poroshell 122 EC-C18 1.9 μm, 2.1x100 mm) at a flow rate of 0.4 mL/min in 5 mM ammonium acetate, pH 5.6 and eluted with a gradient of 5-90% methanol over 6 min. The MS was operated under 125 V skimmer voltage, 100 V nozzle voltage, 350°C drying gas. UV absorbance was recorded from 190-400 nm. The data were analyzed using the MassHunter Qualitative Analysis Software (Agilent). CoA, Ac-CoA, Prop-CoA and Mal-CoA were quantitated by the area of UV absorbance at 258 nm at specific retention time (Figure S7). For MeMal-CoA, the data were analyzed by extracted ion counts (EIC).

To assay acyl transfer, reaction mixtures (20 μL) containing 20 μM EryM, 20 μM EtcB, 1 mM NADPH, 0.1 mM FAD, 1 mM ornithine (Orn) or δ-*N*-hydroxyornithine (hOrn), 1 mM acyl-CoA, 20 mM HEPES pH 8.0 were incubated at 25°C for 1 h. Reactions were quenched and protein precipitated by addition of 60 μL methanol. After centrifuging at 14,000 rpm for 10 min, reaction mixtures were subjected to analysis by LC/MS using an Agilent Q-TOF 6545. For the detection of Orn derivatives, 0.5 μL of each quenched reaction mixture was loaded onto an HPLC column (Agilent Poroshell 120 HILIC-Z 2.7 μm, 2.1x100 mm) at a flow rate of 0.4 mL/min in 0.2% formic acid and eluted with a gradient of 5-95% acetonitrile over 10 min.

### Protein Crystallization and Structure Determination

EryM was crystallized at 4°C by sitting drop vapor diffusion from a 1:1 mixture (4 μL total) of protein stock (10 mg/mL in buffer B) and reservoir solution (1.6-2.0 M (NH_4_)_2_SO_4_, 2-4% (w/v) 2-methyl-2,4-pentanediol, 0.2 M NaCl, 0.1 M sodium cacodylate pH 5.8). Crystals of tetragonal morphology grew over 3-7 days. Crystals were flash-cooled in liquid nitrogen without further cryo-protection. For the Ac-CoA complex of EryM, crystals were soaked in reservoir solution with 8 mM Ac-CoA for 1 h prior to harvesting. SeMet-EryM crystals were grown identically to the wild type. Diffraction data were collected at the Advanced Photon Source (APS) beamline 23-ID-B (EryM free enzyme and Ac-CoA complex) and 23-ID-D (SeMet EryM) (GM/CA@APS) and processed using XDS^50^. The single-wavelength anomalous diffraction (SAD) data for SeMet EryM were analyzed by Phenix^51^ HYSS, AutoSol and AutoBuild to identify Se sites and calculate phase and an electron density map. During refinement, we discovered a cloning error that introduced a Phe substitution for EryM Leu158. The deposited Ac-CoA complex includes Phe158. We corrected the expression plasmid, purified wild-type EryM, confirmed that the substitution had no impact on activity, and collected a new dataset. The deposited structure for the EryM free enzyme includes the Leu158 correction.

EtcB was crystallized at 20°C by sitting drop vapor diffusion from a 1:1 mixture (4 μL total) of protein stock (10 mg/mL in buffer B) and reservoir solution (4-19% PEG 4000, 0.1-0.25 M MgCl_2_, 0.1 M MES, pH 6.5). Crystals of rectangular morphology overnight. Crystals were flash-cooled in liquid nitrogen without further cryo-protection. Diffraction data were collected at the APS beamline 23-ID-B and processed using XDS^50^. The structure was solved by molecular replacement from an AlphaFold2-predicted model^52^. FAD co-purified with EtcB and is present in the refined model at partial occupancy. A large helical lid would cover the active site upon of substrate (hOrn) and cofactors (FAD, NADPH). The lid is disordered and only the ADP moiety of FAD is visible in electron density.

Models were refined using Phenix^51^ with model building in Coot^53^. Structure validation was done with MolProbity^54^ and. The DALI server^55^ was used to search for structure homologs.

### Sequence Alignment and Similarity Network

Sequences were aligned with Clustal^56^. The full-length sequence of EryM was used as a query in a BLAST search against the RefSeq non-redundant database^29^, yielding 5000 hits (maximum). A custom script^30^ was then applied to extract sequences containing both EryM domains (longer than 300 amino acids), maintaining alignment to both EryM domains, resulting in 58 sequences, including EryM.

### Quantification and Statistical Analysis

All biochemical assays were performed in triplicate. Error bars present the standard deviation of the mean.

